# Developmental changes in the morphology and three-dimensional arrangement of antennal hair plates in crickets

**DOI:** 10.64898/2025.12.10.693584

**Authors:** Hui Lyu, Hiroto Ogawa

## Abstract

Antennae of insects are essential mechanosensory organs that facilitate active tactile exploration and spatial navigation. Hair-plate sensilla at the base of the antenna flagellum provide proprioceptive inputs to detect their position. In hemimetabolous insects, such as crickets, the first instar immediately after hatching also possesses antennae, but the developmental dynamics and spatial organization of antennal hair plates remain poorly understood. Here, we present a comprehensive three-dimensional analysis of the antennal hair plates in crickets across developmental stages, from the first instar to adulthood. Using scanning electron microscopy and confocal laser scanning microscopy, we demonstrated that hair plate sensilla were present from the first instars and maintained a highly stereotyped spatial arrangement throughout development. Three-dimensional quantification revealed that new sensilla added during molting were formed at specific sites within the hair plate clusters that had existed at the previous stage, maintaining the spatial pattern despite the substantial growth of the antenna. Multidimensional analyses indicated that the spatial arrangement of sensilla was consistent across individuals, suggesting that organization was genetically determined. Retrograde labeling of sensory afferents showed that sensory neurons in the hair plates converged their axons, extended axon collaterals into the ipsilateral region of the antennal mechanosensory and motor center, and ultimately projected to the subesophageal ganglion. There was no apparent difference in their projection site among hair plates, suggesting no evidence of topographic organization. Our findings highlight the conserved spatial organization of hair-plate sensilla in crickets, suggesting a robust proprioceptive system that provides reliable feedback of antennal position throughout development.

## Introduction

When visual perception is limited, such as in dark or enclosed spaces, many animals rely on active sensing to navigate their surroundings and detect obstacles. For insects, active tactile exploration using the antennae plays a crucial role in spatial perception independent of the visual system. Antennal mechano-sensing mediates object detection and route guidance during obstacle negotiation behaviors in cockroaches (Harley et al., 2009) and stick insects (Krause and Dürr, 2012). In crickets, antennal mechanosensory inputs modulate cercal-mediated escape behavior depending on the shape and position of obstacles, indicating that antennal inputs provide spatial cues that can be used for behaviors elicited by other sensory systems, although they do not necessarily trigger those behaviors directly (Ifere et al., 2022, 2025).

In the insect antennal system, the basal segments are actively moved by muscles at the scape-pedicel (SP) and head-scape (HS) joints, whereas the flagellum lacks intrinsic musculature (Honegger et al. 1990). Mechanosensory receptors located on the scape and pedicel predominantly function as proprioceptors, which provide sensory feedback about antennal position and movements (Toh, 1981). Tactile stimulation of the antennal flagellum elicits the escape response in cockroaches, which is also triggered even if the flagellum is replaced with an artificial object. (Comer et al., 2003). This means that the basal segments of the antenna provide not only self-generated feedback signals but also sensory inputs generated by external stimuli. Thus, the function and organization of mechanoreceptors at the antennal base need to be investigated for understanding the neural substrate of antennal active sensing.

Abundant exteroceptive bristles are distributed at the base of the antenna, as well as proprioceptors, such as the internal Johnston’s organ (JO), external campaniform sensilla (CS), and hair plates (also referred to as Böhm’s bristles) (Staudacher et al., 2005). JO is located beneath the cuticle at the flagellum–pedicel junction and responds to vibratory stimuli, triggering escape responses (Comer and Baba, 2011). CS, arranged in a circular pattern on the distal pedicel, detect cuticular strain and deformation generated at the flexible connection between the pedicel and flagellum (Pringle, 1938; Toh, 1981). Hair plates are distributed on the surfaces of both the scape and pedicel and are thought to be principal proprioceptive sensors precisely detecting antennal position (Okada and Toh, 2000, 2001).

The development and molting-associated renewal of mechanoreceptors have been documented in various hemimetabolous insects, including cockroaches (Moran, 1971; Moran et al., 1976; Schafer and Sanchez, 1973; Schafer, 1997), stink bugs (Brézot et al., 1996), silverfish and bristletails (Thomann et al., 1997), mantids (Carle et al., 2014), and stick insects (Strauß, 2024). In crickets, the developmental processes of mechanoreceptors have been well studied in the cercal organ, where various types of mechanoreceptors are distributed (Gnatzy and Schmidt, 1971; Gnatzy, 1978). During each molting, the old cuticle detaches from the epidermis, and a new cuticle, including sensory structures, forms beneath it (Gnatzy and Römer, 1980). The ventromedial array of cercal clavate hairs, which detects gravity acceleration, shows a consistent spatial position of each sensillum. The sequential addition of sensilla across instars yields a highly stereotyped pattern, enabling each receptor to be individually identified based on its position (Murphey et al., 1980). In contrast to the cercal sensilla, the developmental trajectory of the spatial organization of antennal hair plates remains unknown. Could the precise arrangement of hair plate sensilla required for effective active sensing be established during development?

In addition, afferent pathways of the antennal hair plates have not been identified. No direct projections from hair plates to descending interneurons involved in escape behavior have been found in stick insects (Jaske et al., 2021). It is assumed that mechanosensory inputs from antennal hair plates are initially processed in the central complex, which is involved in spatial orientation and memory, and then activate the motor pathways for escape behavior (Honkanen et al., 2019). The sensory afferents of the cercal filiform sensilla topographically project into the terminal abdominal ganglion, representing the direction of airflow detected by cerci (Paydar et al. 1999). The afferents of antennal hair plates possibly terminate at different sites depending on their location and the antennal joint movements they detect.

To address these questions, we used the field cricket *Gryllus bimaculatus* as a model animal. In this species, dorsoventral movement of the antennae is achieved through the horizontal axis of the HS joint, whereas mediolateral movement is controlled by the vertical axis of the SP joint. These two axes form an orthogonal articulation that allows basic but effective control of the antennal direction (Staudacher et al., 2005). This organization means that crickets’ antennal kinematics are simpler than those in cockroaches and stick insects (Okada and Toh, 2004; Kraus and Dürr, 2004).

In this study, we examined the spatial distribution and fine structure of the hair plate sensilla at the antennal base in crickets, including their developmental changes from the first instar to adulthood, by combining scanning electron microscopy with confocal 3D reconstruction. In addition, retrograde staining of sensory afferents revealed central projections of hair plate sensilla in the cephalic ganglia. Our findings provide important insights into the developmental and anatomical foundations of antennal active sensing, laying the groundwork for future studies on sensorimotor integration in insects.

## Materials and Methods

### Animals

A wild-type strain of crickets (*Gryllus bimaculatus* DeGeer, Hokudai WT strain; Watanabe et al., 2018) reared in our laboratory was used in all experiments. All animals were reared in a 12:12 h light/dark cycle at a constant temperature of 27 °C throughout development. Crickets at various stages, which were nymphal instars N1–N8 and adults <1 week post-imaginal molting, were used to investigate developmental changes in mechanosensory hair plates. The developmental stage of nymphs was determined based on external morphological features, such as body size and wing pad development, as described in a previous study (Suzuki and Nishimura, 1997). Only individuals with intact and undamaged antennae were chosen for analysis to ensure morphological measurements of antennal hair plates. From the sixth instar onward, when sex could be reliably identified by the presence of female ovipositors, only male crickets were used for all experiments. The guidelines of the Institutional Animal Care and Use Committee of the National University Corporation, Hokkaido University, Japan, do not specify any specific treatment requirements for insects in experiments.

### Scanning electron microscopy

Crickets were anesthetized by cooling on crushed ice and then decapitated. The removed heads were soaked in 50% acetone and cleaned using an ultrasonicator (Iuchi US-1; SND, Japan) for 15 minutes. Subsequently, the specimens were dehydrated in an ascending series of acetone solutions (50%, 70%, 90%, and 100%; 5 min each), air-dried, and coated with gold using an ion sputterer (E-1045; Hitachi, Japan). The pretreated specimens were imaged using a field-emission scanning electron microscope (S-3000N; Hitachi, Tokyo, Japan) at an accelerating voltage of 5–10 kV.

### Morphological measurements of the antennal basal segment

The number and length of mechanosensilla on the hair plate were measured by using a confocal microscope (LSM980, Zeiss, Oberkochen, Germany). All tissues except the cuticle were removed using the following treatment: The basal segment of the antenna was isolated from the head and immersed in 1 N NaOH at 60 °C for 24 hours. Specimens were dehydrated through a graded series of 50–100% ethanol, transferred to methyl salicylate for clearing, and placed in a drop of Bioleit reagent (Okenshoji, Tokyo, Japan) on a glass slide. To detect the autofluorescence in the green-to-yellow color range of the cuticle, the basal segment was illuminated with a 488-nm wavelength laser line. Optical section images with a resolution of 1024 × 1024 pixels (pixel size 0.83 μm × 0.83 μm) were captured at 1-µm Z-steps using a confocal microscope equipped with a 10× dry objective (Plan-Apochromat 10×/0.45 M27, Zeiss). Multiple images for one specimen were stitched seamlessly using *ImageJ* (version 1.54; Schneider et al., 2012) to create a three-dimensional reconstruction of the antennal base.

Morphometric parameters of the hair plate, including the length and number of individual sensilla, were measured from the reconstructed images. At all developmental stages (N1–N8 and adults), these morphometric parameters and the spatial distribution of the sensilla were determined for four different positions of hair plates (dorsal, ventral, medial, and lateral) (n = 3 individuals for each stage).

### Data processing

Three-dimensional locations of individual hair plate sensilla were determined based on the confocal images and organized for each specimen. For spatial alignment and dimensionality reduction of the morphological parameters, principal component analysis (PCA) was used in R (version 4.5.0; R Core Team, 2025). Three-dimensional coordinates for each sensillum were centered by subtracting the mean position of all sensilla in each hair plate, and then PCA was performed using the built-in prcomp function to derive the major axes of spatial variation. The coordinate data of the sensilla were projected onto these principal axes, which were X_pca, Y_pca, and Z_pca, enabling comparison of the sensilla distribution pattern across different samples and stages. Spatial similarity in the distribution of sensilla between samples was quantified using pairwise Hausdorff distances (Alt et al., 2003) computed with the NumPy (Harris et al., 2020) and SciPy (Virtanen et al., 2020) libraries in Python (version 3.13).

### Anterograde staining of sensory afferents

Adult male crickets were used for anterograde staining. Crickets were anesthetized on crushed ice to prevent their movements. After cutting the sensilla of one or two hair plates on the scape or pedicel with a razor blade, a glass micropipette filled with a dye solution containing 1% Alexa Fluor 488 or 1% Alexa Fluor 594 dextran conjugates (10,000 MW; Thermo Fisher Scientific) diluted in distilled water was placed in contact with the cut area. To perform double staining of multiple hair plates, two micropipettes, each filled separately with Alexa Fluor 488 and 594, were placed in contact with different target regions. The glass pipettes were secured with clay, and the specimens were kept in a moist chamber at 4 °C overnight.

The cephalic ganglia, including the brain, suboesophageal ganglion, and ventral nerve cords connecting them, were then isolated, transferred onto filter paper for shaping, and fixed with 4% paraformaldehyde (PFA) for 6 hours. After fixation, the ganglia were dehydrated in an ascending ethanol series (50%, 70%, 90%, and 100%; 5 min each) and then cleared in methyl salicylate.

### Confocal imaging of sensory afferents

The stained mechanosensory afferents of the hair plates were imaged using a confocal laser scanning microscope (LSM980, Zeiss) equipped with a 10× dry objective (Plan-Apochromat 10×/0.45 M27, Zeiss). Optical sectioned images with a resolution of 1024 × 1024 pixels (pixel size 0.83 μm × 0.83 μm) were captured at 1 μm Z-steps from the ventral to dorsal side of the ganglion. Using *ImageJ* (version 1.54), the Z-stack series of images was merged into a single image and adjusted for brightness and contrast to clearly visualize the full morphology of the afferents.

## Results

### Morphological features of the hair plates at the antenna base segments

Scanning electron microscopy (SEM) revealed a detailed structural organization of the cricket antenna base. The basal segments comprise the scape and pedicel, which serve as the foundation for antennal movements (Fig. 1a). High-magnification images showed the hair plate sensilla in the scape, each of which was embedded within a flexible socket structure (Fig. 1b; Toh Y, 1981). The sensilla were located in direct contact with the articular membrane at the basal joints (Fig. 1c), indicating that they could be deflected by adjacent membranes when the antennal segments moved relative to each other. Hair plates were already present at the first instar stage, along with circularly arranged CS and bristle hair sensilla (Fig. 1d). SEM observations at the second instar, fourth instar, and adult stages revealed that the sensilla of the ventral hair plates had a similar and stereotyped spatial arrangement, which appeared to be maintained throughout development (Fig. 1e‒g). The position of the hair plates gradually shifted toward the proximal end of the scape as the developmental stage progressed. This positional shift might indicate a developmental adaptation that helps these hair plates precisely detect the movements of antennal segments, even as body size increases.

**Fig. 1.**
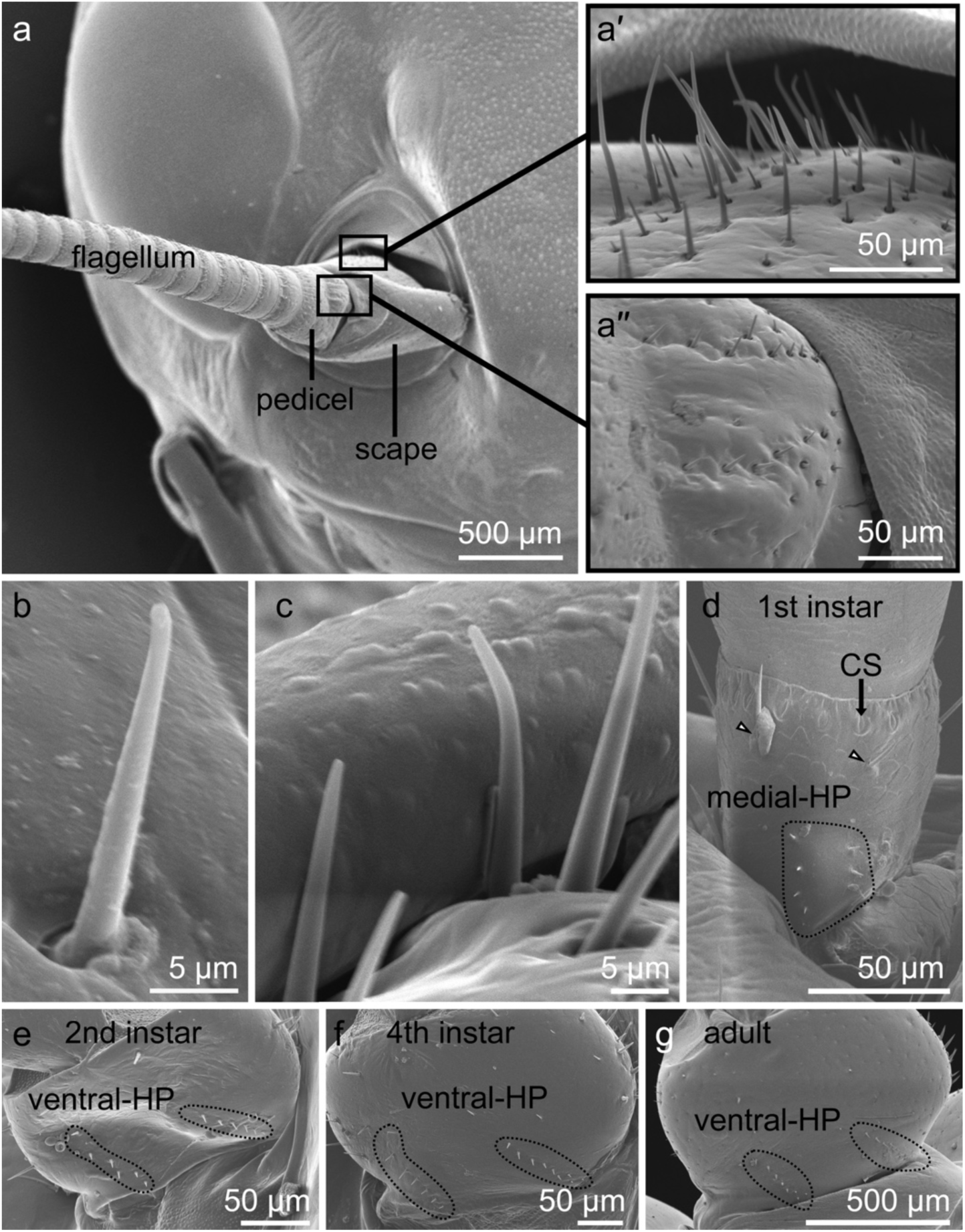
Scanning electron micrographs of the antennal base segments of *Gryllus bimaculatus*. **a** Frontal view of the cricket head showing the scape, pedicel, and filamentous flagellum. Higher-magnification views of the hair plates on the scape (**a′**) and pedicel (**a″**). **b** A single hair plate sensillum on the scape, embedded in a flexible socket at the joint surface. **c** Hair plate sensilla deflected by the adjacent joint membrane. **d** Medial view of the antennal base in a first instar nymph. White arrows indicate exteroceptive bristles. **e–g** Ventral-HP viewed from the ventral side of the scape in the (**e**) second instar, (**f**) fourth instar, and (**g**) adult. HP, hair plate; CS, campaniform sensilla.

### Three-dimensional distribution of the hair plate

To further investigate the detailed spatial organization of hair plates, we created three-dimensional reconstructions of the antennal base across various developmental stages, based on confocal microscopic images (Fig. 2). Four distinct groups of hair plates were consistently observed across individuals and developmental stages: the dorsal, ventral, medial, and lateral groups. The dorsal and ventral hair plates are located on the posterior surface of the scape, which were thought to detect dorsoventral movements (elevation) of the antenna. The sensilla of the dorsal hair plate were arranged linearly in one to several rows adjacent to each other in the proximal-distal direction, while the sensilla of the ventral hair plate were arranged in one or two rows separated from each other (Figs. S1, S2). Meanwhile, the medial and lateral hair plates are located on the posterior surface of the pedicel, which likely detect mediolateral (horizontal) movements of the antenna. These hair sensilla were also arranged in a few rows, from proximal to distal (Figs. S3, S4). These stereotyped and region-specific arrangements of the hair plate sensilla on the scape and pedicel were already observed at the first instar, suggesting that this spatial distribution was conserved throughout development.

**Fig. 2.**
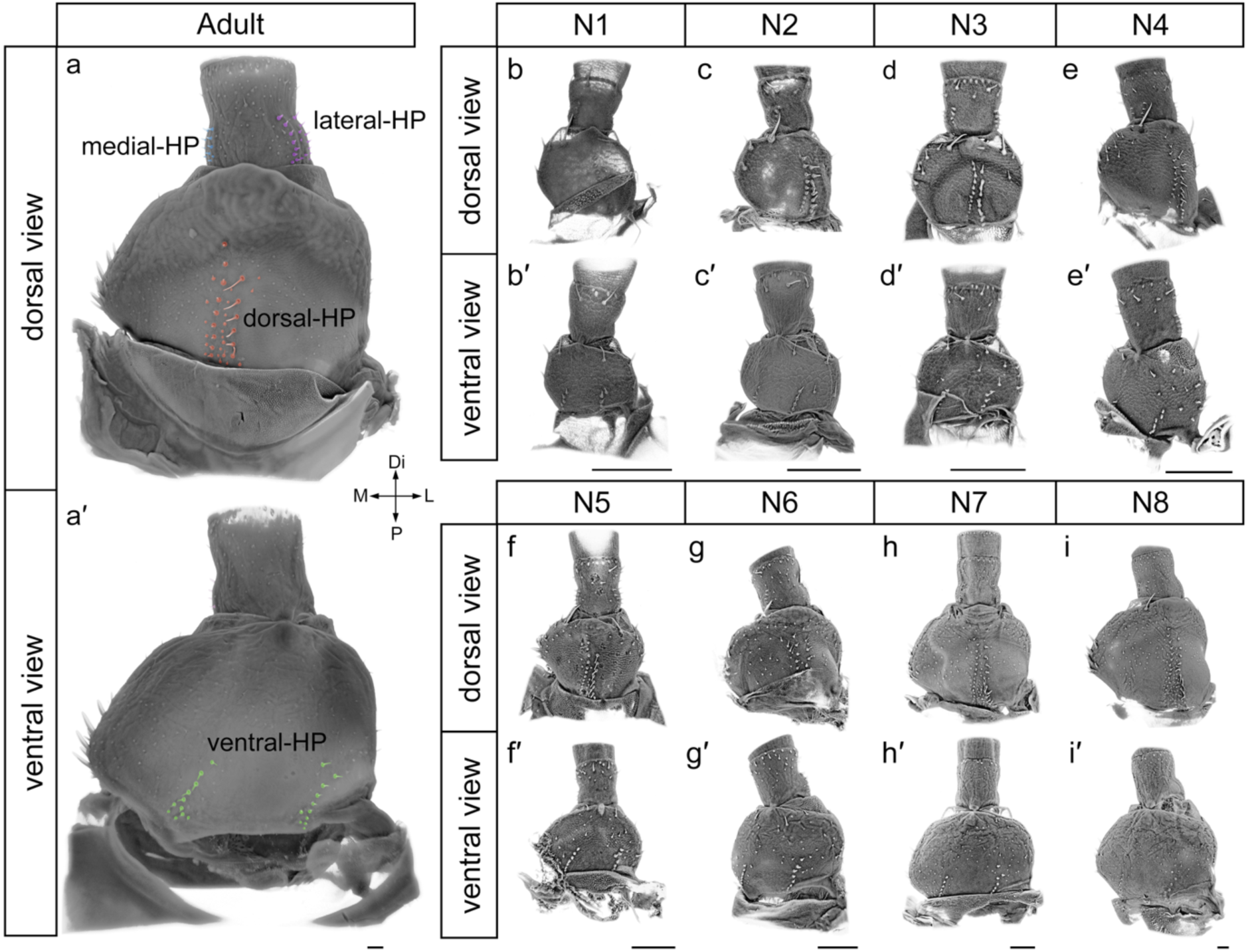
Three-dimensionally reconstructed antennal base and spatial distribution of hair plates across developmental stages. **a** Dorsal (**a**) and ventral (**a′**) views of the antennal base in adult crickets. Sensilla of the medial, lateral, dorsal, and ventral hair plates were colored in cyan, purple, orange, and green, respectively. **b–i** Dorsal (**b–i**) and ventral (**b′–i′**) views of the antennal base at different nymphal stages, N1 to N8. Scale bars: 100 μm.

Moreover, to quantify developmental changes in the arrangement of sensilla as the antennal base segments grew, we compared the spatial patterns of their locations by normalizing the hair plate size (Fig. S1‒S4). As the developmental stage progressed, the number of sensilla for each hair plate increased. By adding new sensilla at specific locations, the sensilla cluster of each hair plate expanded without altering the overall spatial pattern. The sensilla within each hair plate were distributed in a compact, constrained area, and their relative positions appeared to stay consistent across developmental stages. Even as the number of sensilla increased, their arrangement maintained the original shape and direction for each plate. In the dorsal hair plate, two distinct rows of sensilla were observed from the N1 to N3 instars. In the N4 instar, the third row of sensilla began to appear between the previous two rows, and in the N5 instar, this row became more prominent. In the N6 instar, a fourth row of sensilla appeared along the medial edge of the dorsal hair plate, toward the midline of the antenna. In contrast, the ventral hair plate maintained a two-row layout throughout development, although in the instars older than the N6 stage, additional short sensilla appeared close to the proximal region. The medial and lateral hair plates exhibited almost symmetrical and similar arrangements of sensilla. In instars older than the N6 stage, a third row of shorter sensilla developed between the previous two rows on these hair plates.

This three-dimensional organization conserved across development suggests that hair plate placements are genetically regulated and that the mechanosensory system maintains its functional topology despite substantial size increases during maturation. The alignment of hair plate positions with the major axes of antennal movements is effective for detecting specific directional displacements, suggesting their roles in proprioceptive feedback in cricket antennae.

### Developmental dynamics and morphometrics of antennal basal segments

To elucidate developmental changes in antennal hair plates, we examined their morphometric parameters, such as the size and number of sensilla in four regions of the basal segments, across different developmental stages. For quantitative analysis of the development in basal segment size, we measured the straight-line distance between the proximal and distal ends of each segment, based on 3D confocal image data. Both the scape and pedicel rapidly increased in the proximal-distal length throughout development, from the first instar (N1) to the adult stage (Fig. 3b). Notably, the segment length showed a significant increase from the fifth to seventh instars (N5–N7). On the other hand, the nearest-neighbor distances between sensilla increased almost linearly in all four regions of the hair plate throughout development (Fig. 3c). This means that the expansion in inter-sensilla space was relatively smaller than the overall enlargement of the hair plate surface area with maturation. For example, the pedicel increased in length by up to fivefold from N1 instar to adult (mean length: 113.0 ± 3.0 μm in N1, 591.3 ± 23.6 μm in adults), and the scape showed an even greater expansion of eightfold (mean length: 140.2 ± 4.5 μm in N1, 1172.3 ± 78.1 μm in adults). In contrast, the distance between nearest sensilla increased by about 2.5-fold (mean distance: 25.1 ± 0.5 µm for the dorsal hair plate, 23.0 ± 0.8 µm for the lateral hair plate, respectively). The increasing rate of inter-sensilla distance was nearly comparable among the dorsal, ventral, medial, and lateral hair plates, indicating that they continued to share the same spatial scale during development.

**Fig. 3.**
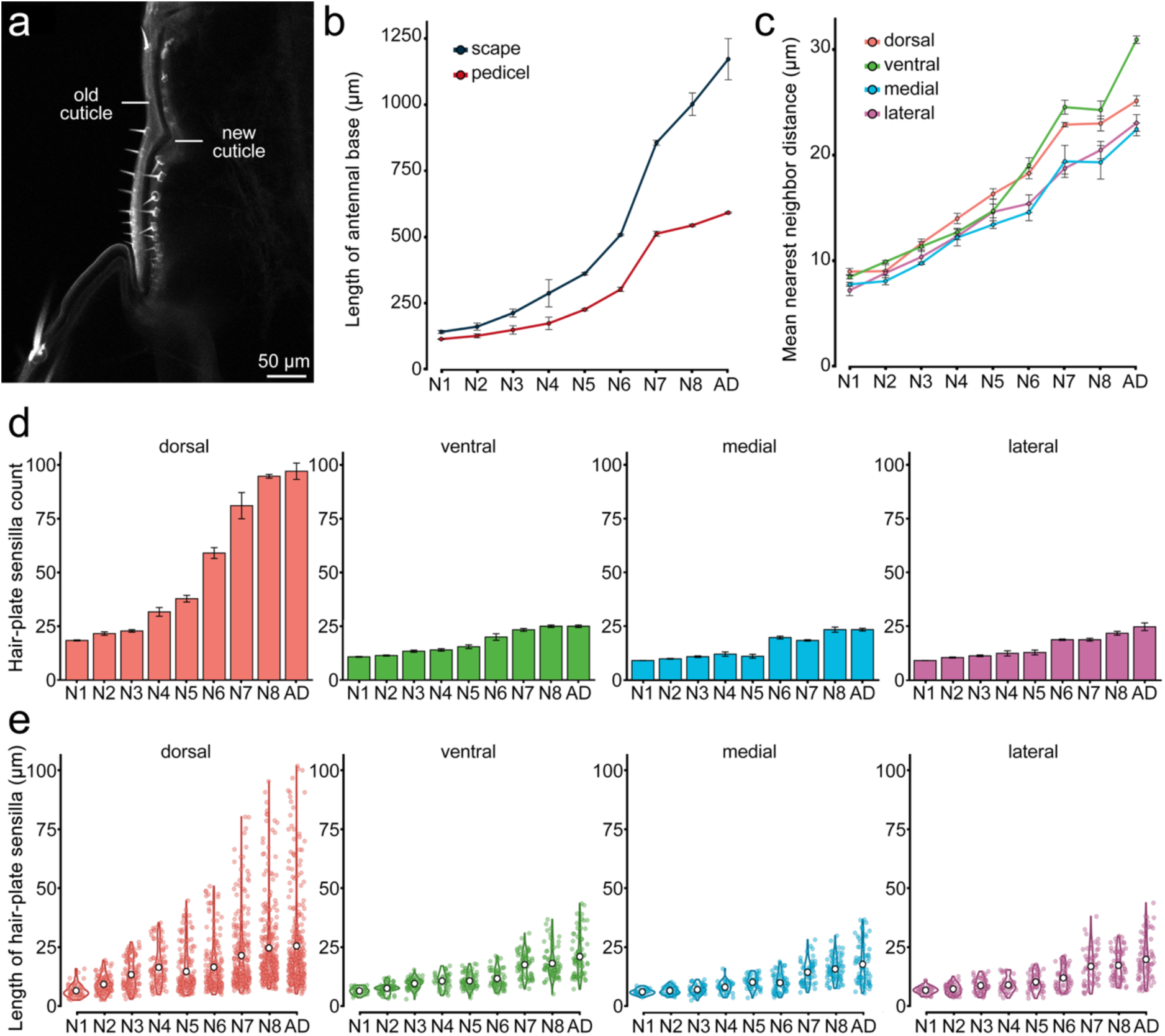
Developmental changes of antennal basal segments and hair plate sensilla. **a** A confocal image of the antenna during the molting from the final nymphal stage (N8) to adulthood. **b, c** Developmental curves of the antennal base segments (scape and pedicel, **b**) and hair plates (**c**) across developmental stages (N1–N8, AD: adult). The values for the vertical axis in **b** and **c** indicate the mean of the proximal-to-distal length of base segments (**b**) and the mean distance between the nearest sensilla within different locations of hair plates (**c**). Error bars indicate ± SEM. **d, e** Mean number of sensilla (**d**) and the length of individual sensillum (e) within the dorsal, ventral, medial, and lateral hair plates at different developmental stages. Error bars in (**d**) represent ± SEM. Colored open circles represent the data for each sensillum, and white filled circles indicate the median values.

Although new sensilla were added locally during development, the overall spacing between them still increased. These findings suggest that the addition of sensilla partially compensates for the reduction in spatial resolution caused by segmental growth. The number of sensilla increased progressively throughout development for all four locations of the hair plate. The dorsal hair plate in particular had more sensilla than the others, and this difference increased as the developmental stage progressed (Fig. 3d: The number of sensilla in adults was 97.0 ± 3.8 for the dorsal, 25.0 ± 0.6 for the ventral, 23.3 ± 0.7 for the medial, and 24.7 ± 1.8 for the lateral hair plates, respectively). Alongside the increase in sensilla counts, the hair length of individual sensilla also elongated and became varied as the cricket developed (Fig. 3e). The dorsal hair plate sensilla showed the greatest variation in their length. Even as the hair length became more varied, shorter sensilla remained to account for the majority across the developmental stages. This result suggests that the hair sensilla from the previous stage are replaced by elongated ones, with new, short sensilla added simultaneously during molting. These developmental changes in the existing areas, number, and external morphology of the antennal hair plates are possibly involved in maintaining and refining their proprioceptive function in the active sensing of the antennae.

### Multidimensional spatial analysis of hair plate sensilla

To quantitatively assess the consistency of hair plate spatial organization throughout development, we compared the distribution patterns of individual sensilla across samples and developmental stages using the Hausdorff distance of their three-dimensional coordinates (see Methods). This metric, indicating the maximum spatial deviation between the nearest two sensillar sensilla, can be used as a measure of geometric dissimilarity. We analyzed this metric separately for the dorsal, ventral, medial, and lateral hair plates and visualized pairwise distances between samples at the same and different developmental stages, as a heatmap (Fig. 4).

**Fig. 4.**
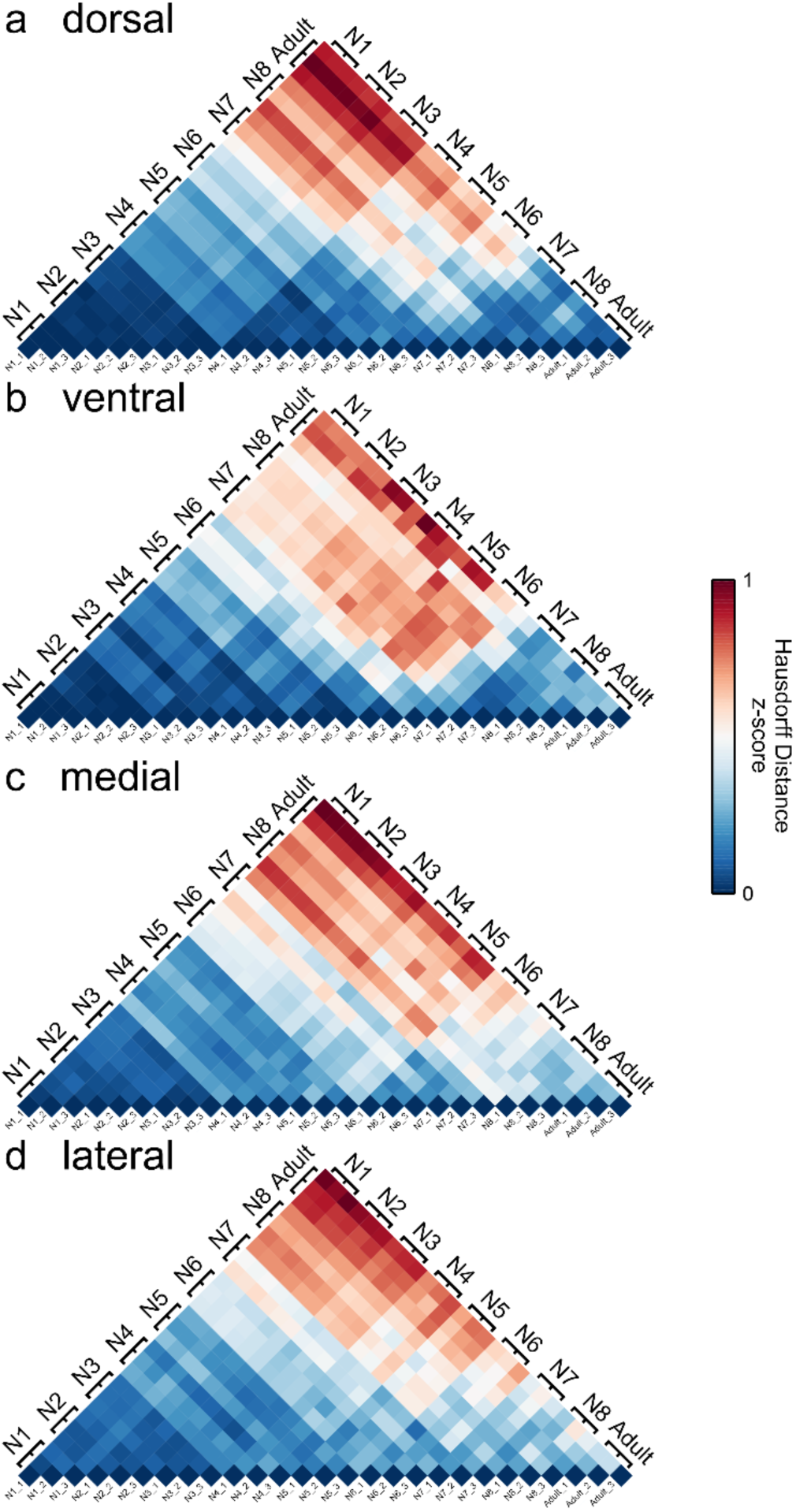
Similarity in spatial arrangement of hair plate sensilla across developmental stages **a–d** Heatmaps depicting pairwise spatial similarity in the spatial distribution of hair plate sensilla among samples for four different locations: dorsal (**a**), ventral (**b**), medial (**c**), and lateral (**d**). The colors for each matrix entry represent the normalized Hausdorff distance (z-score) between two samples calculated from the three-dimensional arrangement of the hair plate sensilla. As shown in the color table on the right, blue represents a shorter distance, indicating greater spatial similarity, while red represents a longer distance, indicating lower similarity.

The Hausdorff distances for all four hair plates were smaller between samples at the same developmental stage than those at different stages, suggesting that the distribution pattern of hair-plate sensilla was maintained reproducibly across individuals. Regarding the inter-stage variabilities among the four hair plates, the dorsal plate showed the smallest distances, indicating that the spatial arrangement of sensilla was consistent, especially in the dorsal plate throughout development. The sensillar distributions aligned by the PCA method revealed that the number of sensilla increased by adding new sensilla to fill gaps between those existing at the previous stage (Fig. S1–S4). In addition, the length of individual sensilla on the hair plate correlated with their position within the hair array, with longer sensilla located distally than proximally. Furthermore, on the dorsal hair plate, hair length increased from the medial toward the lateral sides (Fig. S1). These overall spatial patterns of sensilla remained largely unchanged throughout developmental stages, without displacement or clustering shifts. Together, these results demonstrated that the spatial architecture of the hair plate system was highly conserved throughout development. The particularly low variation in the dorsal hair plate suggests that its spatial sensillar layout was tightly regulated. This possibly reflects its role in monitoring vertical antennal movements. These spatial organizations likely ensure consistent proprioceptive feedback during active sensing as the cricket grows.

### Central projection of hair plate afferents

To examine the central projection of sensory neurons of hair plate sensilla into the cephalic ganglia, including the brain, we performed anterograde labeling from hair plates at different locations. Confocal images of the stained afferents showed that these axons branched to collaterals within the lateral region of the dorsal lobe (DL) in the brain, where their axons further descended through the neck connective to the ipsilateral region of the subesophageal ganglion (SOG) (Fig. 5). The axon collaterals ran dorso-medially and terminated in the dorso-medial region of the DL, which corresponds to the antennal mechanosensory and motor center (AMMC, Yoritsune and Aonuma, 2012). The sensory afferents, including collaterals, terminated within the ipsilateral hemisphere of the ganglia and never extended across the midline. This projection pattern was consistent across all anatomical positions of the hair plates, indicating that the afferent organization was not likely position-specific but shared among dorsal, ventral, medial, and lateral plates (Fig. 5a–e). Although afferent fibers from the dorsal, ventral, medial, and lateral hair plates entered the AMMC through partially segregated routes, their axonal trajectories overlapped substantially within both AMMC and SOG. No clear differences in their terminated regions, suggestive of topographic mapping, were observed. This result suggests that positional information about the antennal movement axis from the periphery is not spatially represented in the processing center but rather decoded by specific postsynaptic neurons, such as the labeled-line system. Anterograde labeling of the sensory afferents also visualized the cell bodies at the base of the antenna, where the axons were originated from spatially dispersed cell bodies and converged into distinct axon bundles (Fig. 5f).

**Fig. 5.**
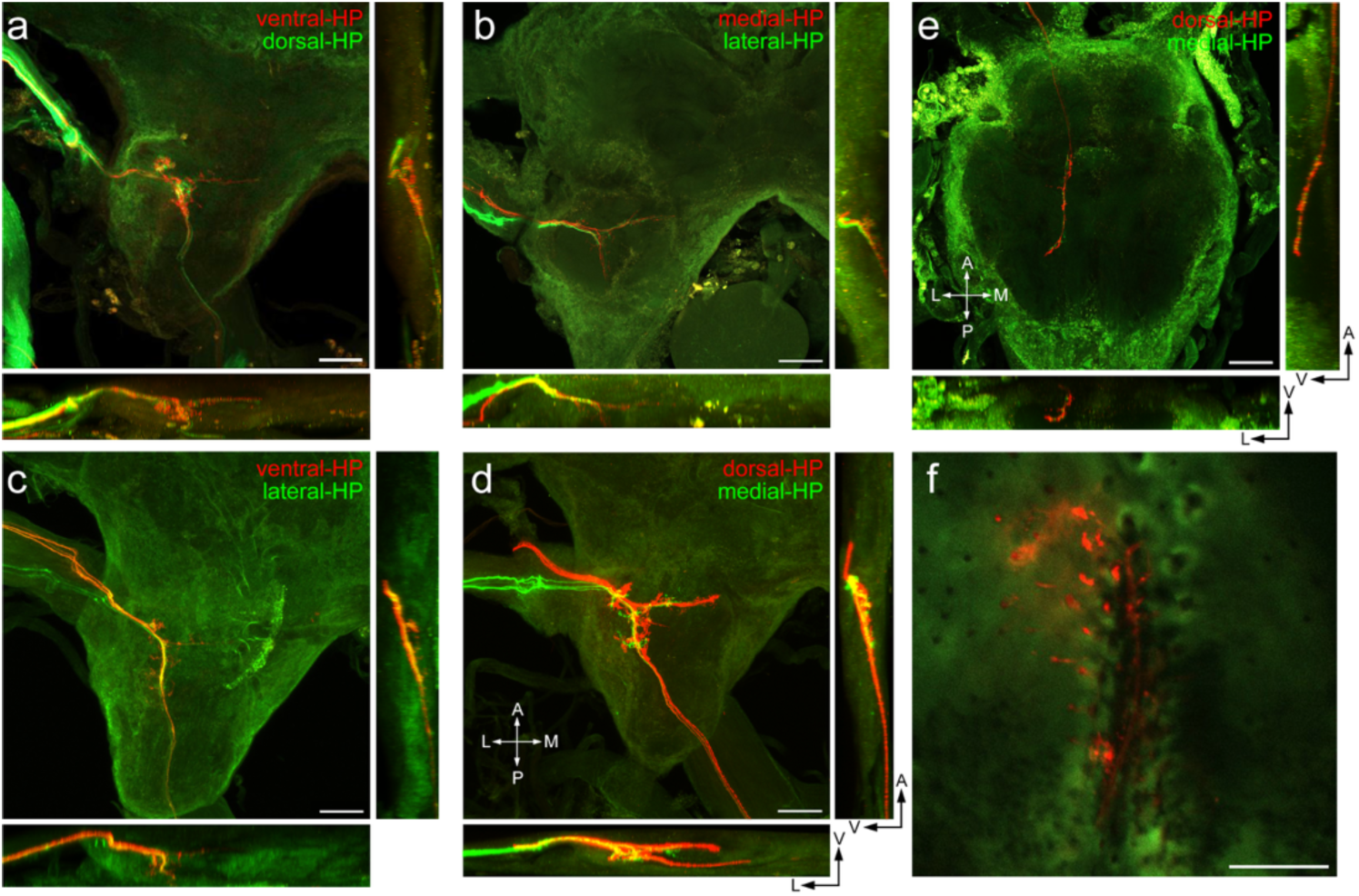
Central projections of mechanosensory afferents from hair plates. **a-d** Fluorescent image of deutocerebrum showing anterograde double labeling of sensory afferents from the ventral and dorsal hair plates (**a**), from the medial and lateral hair plates (**b**), from the ventral and lateral hair plates (**c**), and from dorsal and medial hair plates (**d**). **e** Fluorescent image of SOG showing afferent projections from dorsal and medial hair plates. **f** A confocal stack image showing peripheral branches of sensory afferents for the dorsal hair plate. Cell bodies are observed at the position close to the base of individual hair sockets, and their axons converge into a bundled projection. Sensory neurons are labeled in red, and cuticle autofluorescence is shown in green. Scale bars: 100 μm

## Discussion

### Three-dimensional spatial organization and development

Crickets are valuable model insects for studying the antennal mechanosensory system because their basal antennal joints operate along two orthogonal axes that separately control horizontal and vertical movements (Staudacher et al., 2005). Unlike other insects with more flexible or variably angled joint configurations, the cricket antennae movements follow a relatively simple and stereotyped pattern. Correspondingly, hair plate sensilla are strategically positioned to directly monitor these two major axes of antennal movements. In *Gryllus bimaculatus*, the number of hair plate sensilla exceeds that reported for *Acheta domesticus* (dorsal: 52, ventral: 17, medial and lateral: 15), possibly due to its larger body size (McCarter and Loudon, 2025). The combination of a simple joint structure and consistent sensillar arrangement will provide an accessible system for studying how proprioceptive input is spatially organized and developmentally regulated.

In this study, we systematically mapped the three-dimensional morphology and spatial distribution of hair plate sensilla from the first instar to adulthood (Fig. 2). Our results revealed that the spatial arrangements of hair plate sensilla were highly consistent across individuals and developmental stages, indicating a genetically determined organizational pattern (Fig. 4; Fig. S1–S4). As the cricket developed, existing sensilla became more widely spaced, and new sensilla were added in specific regions, such as between longitudinal rows, along the peripheral margins, or near the base. This non-uniform but structured enlargement preserved the overall arrangement, maintaining the original spatial organization. This conserved organization likely ensures stable proprioceptive inputs even as the animal grows and its antennae elongate. This conserved arrangement of sensory organs may be controlled genetically, as demonstrated for the patterning of mechanosensory bristles in *Drosophila*, where the arrangement of epidermal sensory organs is regulated by genes such as hairy and by self-organized Notch signaling dynamics (Orenic et al., 1993; Corson et al., 2017).

Interestingly, sensilla of distinct lengths were consistently located at corresponding relative positions in the antennal base segment across individuals and developmental stages. This conserved distribution pattern related to hair length suggests that the spatial organization of hair plate sensilla is tightly regulated during development. Such consistency is likely required for their function as proprioceptors, which accurately encode self-generated antennal movements based on the geometric structure of the antennal base. Other insect mechanosensory systems, such as the filiform hair sensilla on the cricket cerci, serve as exteroceptive organs to detect unpredictable mechanical stimuli like air currents. These exteroceptive systems contain large numbers of receptors distributed across broad surfaces (Shimozawa and Kanou, 1984). As shown in quantitative studies on the cercal sensory system, although interindividual variability in the number and placement of hair sensilla is relatively greater, the overall sensillar distribution on the cerci is organized based on their preferred direction of airflow stimuli, which is well approximated by computational models (Miller et al., 2011; Heys et al., 2012). This organized pattern of hair array is thought to enable robust detection by reducing noise through the high density of receptors. In contrast, the antennal hair plate system had fewer sensilla, which were more spatially restricted and aligned precisely with the genetically regulated joint morphology. The consistent spatial pattern observed in all developmental stages likely reflects the proprioceptive roles of these receptors, which require providing reliable and steady inputs throughout postembryonic growth.

### Sensory representation of antennal movements

Neural tracing demonstrated that all hair plate afferents projected to both the AMMC and the SOG, but did not extend into the ventral flagellar afferent (VFA) region. This projection pattern closely parallels recent findings in the cockroach brain, where sensory afferents of the scapus-pedicel joint project densely to both the medial and lateral subregions of the AMMC, but not to the VFA (Althaus et al., 2022). Calcium imaging studies of mechanosensitive descending neurons have shown that tactile stimulation to different locations of the antenna flagellum induces calcium increases in distinct dendritic regions, suggesting topographic representation of the antennal mechanosensory inputs (Bayley and Hedwig, 2018).

Furthermore, histological studies in cockroaches demonstrated that mechanosensory afferents from the antenna flagellum exhibit somatotopic organization, in which sensory neurons located at the more distal site of the flagellum project to progressively more medioventral regions in the deutocerebrum and SOG (Nishino et al., 2005).

In contrast, although projections from different hair plates showed partially distinct collateral patterns, there was no evidence of a clear topographic map within the AMMC or SOG (Fig. 5). These findings suggest that positional information provided by antennal hair plates is transmitted to the specific postsynaptic interneurons and integrated by more central circuit, rather than being mapped in a primary sensory center, as observed for flagellar input. This supports the view that the AMMC functions as an integrative hub to relay multiple mechanosensory inputs and to mediate active antennal movements. In AMMC, the hair plate inputs are considered crucial for proprioceptive feedback during self-generated movements, which differ from those of flagellar afferents providing exteroceptive inputs. In contrast, the cercal sensory system, which is a typical exteroceptive organ, is known as an example of an afferent map, where the mechanosensory afferents of filiform hair sensilla topographically projected into the terminal abdominal ganglion based on their directional sensitivity (Jacobs and Theunissen, 1996; Paydar et al., 1999). Wind-sensitive giant interneurons (GIs) primarily decode the directional information of airflow detected by cerci, depending on the relative positions of their dendrites within the spatial map formed by mechanosensory afferents (Jacobs and Theunissen, 2000; Ogawa et al., 2008). Due to the considerable individual variability in the number of filiform hairs on the cerci (filiform hairs are significantly delicate and can easily be removed in accidents), synaptic integration based on the spatial relationship between afferent inputs and the dendritic architecture of GIs is necessary to achieve reliable coding of airflow dynamics. In the antennal hair plate system, however, the morphological organization of the fewer sensilla is already highly stereotyped and consistent across individuals. This precise arrangement at the sensory receptor level likely ensures that the direction and movement information of the antenna can be reliably transmitted to postsynaptic neurons and central circuits, even without a topographic map at the primary processing stage. Thus, unlike the cercal system, the afferent pathway of the antennal hair plate may rely less on central spatial mapping and more on the innate organization and developmental consistency of its input layer.

### Proprioceptive information provided by the antennal hair plates

Previous physiological studies have shown that scapal hair plate receptors of cockroaches exhibit phasic-tonic discharge in response to antenna deflection (Okada and Toh, 2001). The variety in sensillar morphology, including differences in length and orientation, likely enables the hair plate system to detect a broad range of antennal movement dynamics. The computational model of Ache and Dürr (2015) demonstrated that antennal movement gradually deflects individual sensilla, so that the greater the joint displacement, the more sensilla are activated in the hair plate. This suggests that both the activity of individual sensilla and the population signal reliably encode the joint angle correlated with antennal movements.

However, their model did not include variation in sensillar length. Our morphological data revealed that the hair lengths varied depending on their position within the organized spatial pattern. Specifically, shorter hairs were typically located closer to the joint, where they could be activated by a smaller change in joint angle. In contrast, longer hairs positioned farther from the joint would respond to a greater displacement. This systematic distribution likely provides a functional architecture that enables continuous and fine-scale monitoring of antennal joint angles. Previous behavioral studies have clearly shown that crickets and cockroaches can perceive antennal orientation with high accuracy (Okada and Toh, 2004; 2006; Ifere et al., 2022; 2025). Proprioceptive information about antennal orientation may be encoded through different mechanisms from the topographic map.

### Functional significance of reafferent inputs for active sensing

Many species of insects, such as crickets and cockroaches, actively move their antennae to sense their surroundings. Thus, antennal mechanosensory signals can originate from either self-generated movements or passive contact with external objects. Our findings support the idea that antennal hair plates serve as key proprioceptors that encode antennal movement during active sensing (Ache and Dürr, 2015). Similar mechanisms have been observed in cockroaches, stick insects, and moths, suggesting a conserved role of antennal hair plates across insect species (Okada and Toh, 2004; Jaske et al., 2021; Krishnan et al., 2012). Owing to its robust anatomical organization and genetic consistency, the cricket serves as a powerful model for unraveling the principles of proprioceptive feedback and active sensing.

A recent theory frames active sensing as a closed-loop “sensory-energetic loop” (Ahissar and Assa, 2016). In this framework, alloactive sensing describes situations where an animal adjusts its sensors to detect the environment, whereas homeoactive sensing refers to cases where the animal alters its environment. In the tactile sensory system, both types of sensing often occur simultaneously; therefore, animals must predict the sensory consequences or reafferent inputs generated by their own actions (Zweifel and Hartmann, 2020). Unlike typical active sensing frameworks, however, reafferent inputs mediated by the activation of hair plates during active scanning of antennae do not alter the environment. In this context, rather than predicting sensory outcomes, it is necessary to monitor in real time which area of space the antenna is currently exploring. It is proposed that this information is provided by proprioceptive inputs from the hair plates, enabling spatial awareness through antennal exploration (Ifere et al., 2022; 2025).

The precise and reproducible spatial organization of hair plate sensilla allows the spatially structured recruitment of receptor activation, which may contribute to a reliable neural representation of antennal position. Future physiological studies on postsynaptic neurons downstream of the hair plates will clarify how antennal angles are decoded. In addition to these experiments, the computational model based on the morphometric data of hair plate sensilla, including their three-dimensional distribution and length reported in this study, as well as antennal kinematics, will clarify how reafferent information is represented during active tactile exploration.

## Acknowledgments and Funding Information

### Funding

This work was supported by JSPS KAKENHI Grant No. 21H05295, 21K06259 to H.O. and JST SPRING Grant No. JPMJSP2119 to H.L.

### Author Contributions

H.L. and H.O. conceived the project and designed the experiments; H.L. performed the experiments; H.L. analyzed the data; H.L. and H.O. wrote the manuscript.

### Data Availability

The datasets generated during the current study are available from the corresponding author on reasonable request.

### Additional information

Competing financial interests: the authors declare no competing financial interests.

