## Supplementary Figures for "Developmental changes in the morphology and three-dimensional arrangement of antennal hair plates in crickets"

**Supplementary Figure 1**

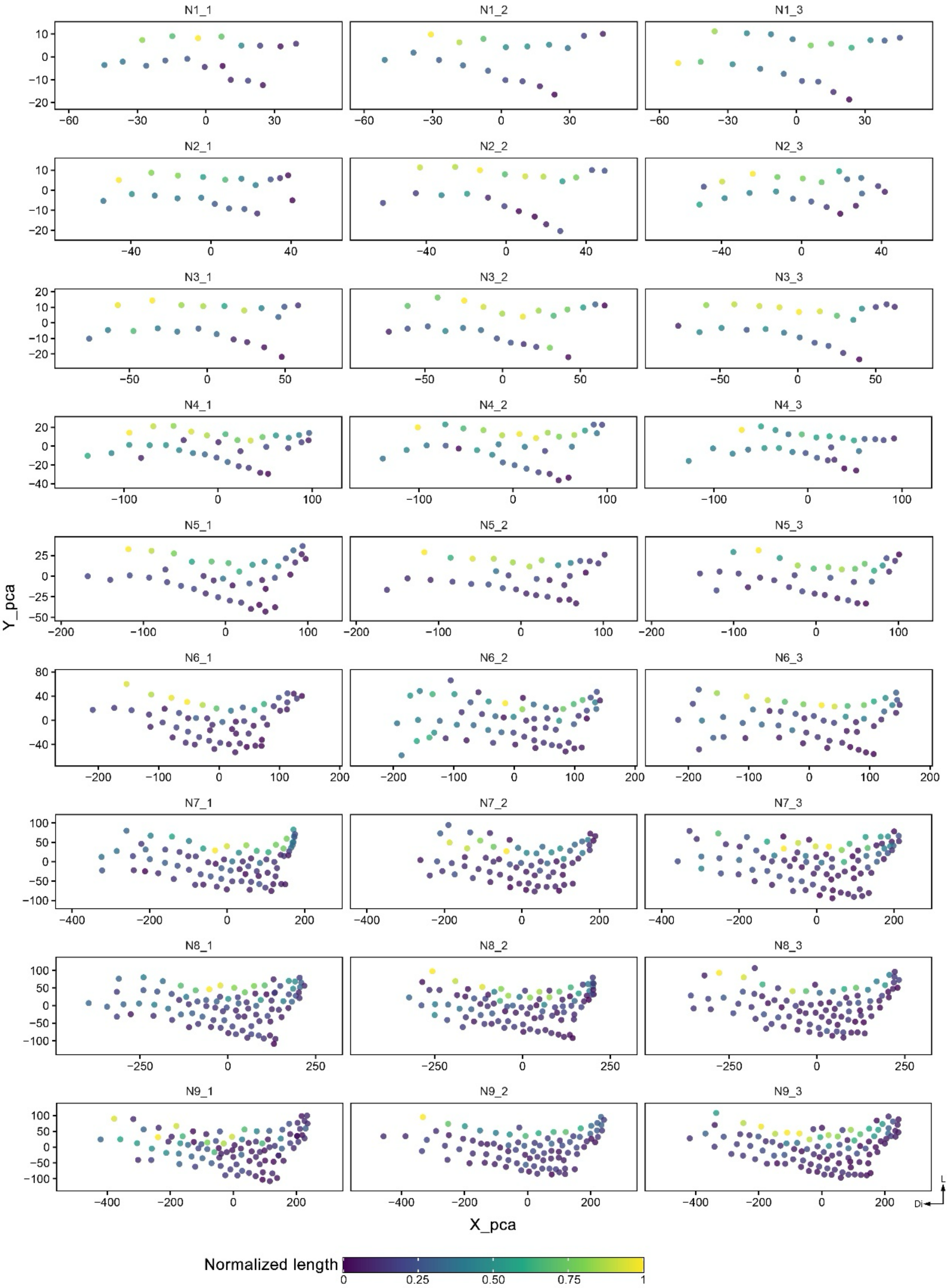


**Fig. S1 Standardized spatial distributions of dorsal hair-plate sensilla across developmental stages.** Each dot represents an individual sensillum for the dorsal hair plate of the antennal base, plotted in the standardized XY-plane after principal component analysis (PCA) alignment (See Material and Methods). The data were obtained from 3 different samples for each developmental stage. Colors of dots indicate the normalized length.

**Supplementary Figure 2**

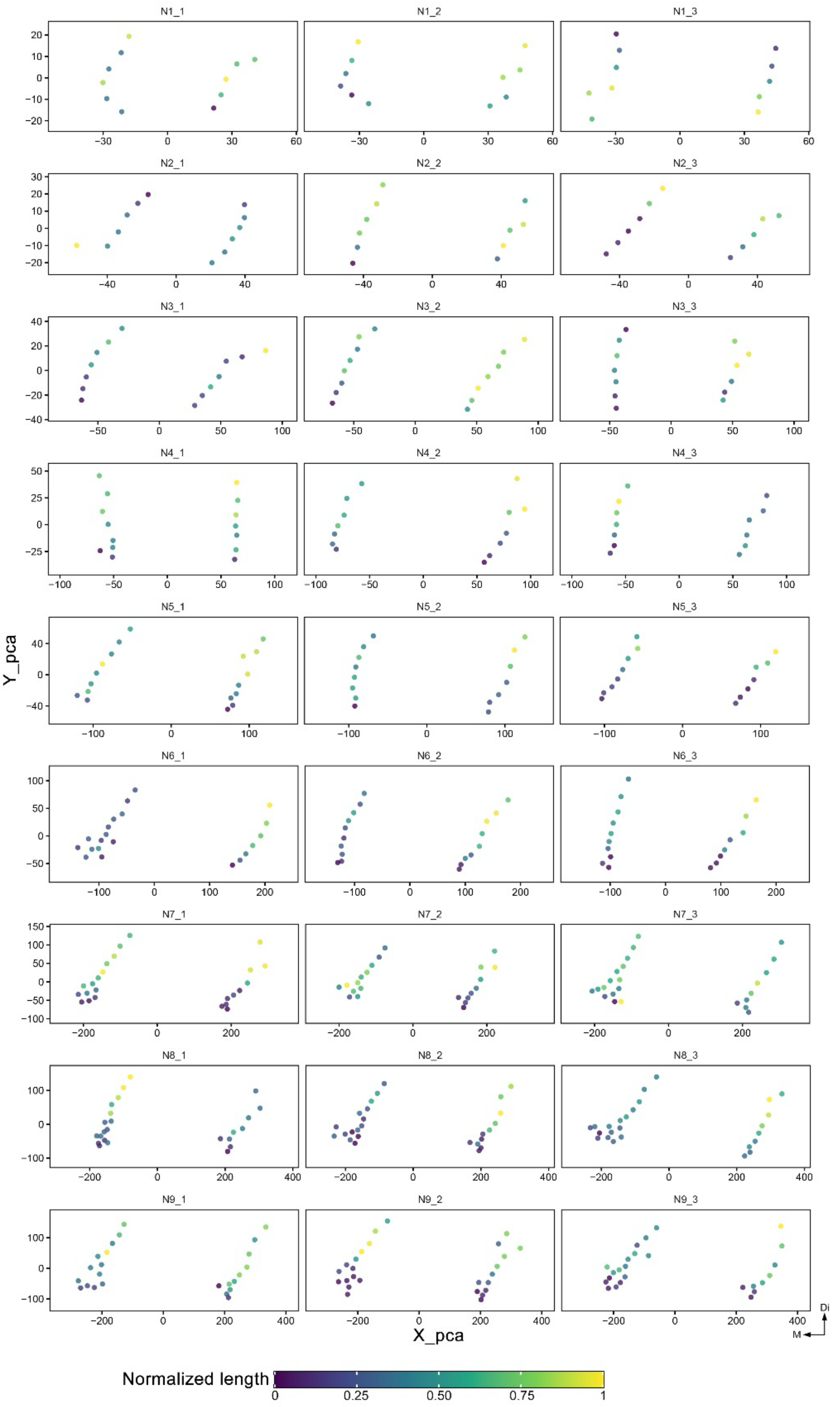


**Fig. S2 Standardized spatial distributions of ventral hair-plate sensilla across developmental stages.** Each dot represents an individual sensillum for the ventral hair plate of the antennal base, plotted in the standardized XY-plane after principal component analysis (PCA) alignment(See Material and Methods). The data were obtained from 3 different samples for each developmental stage. Colors of dots indicate the normalized length.

**Supplementary Figure 2**

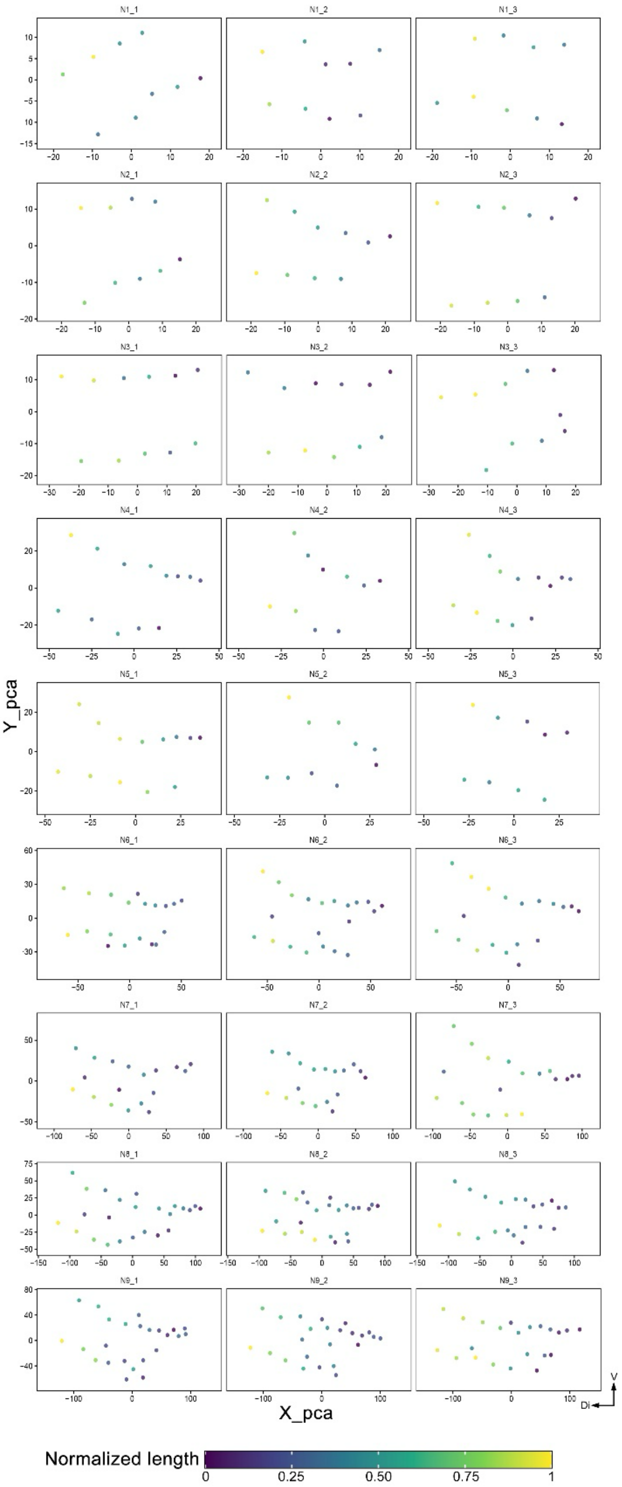


**Fig. S3 Standardized spatial distributions of medial hair-plate sensilla across developmental stages.**

P Each dot represents an individual sensillum for the medial hair plate of the antennal base, plotted in the standardized XY-plane after principal component analysis (PCA) alignment (See Material and Methods). The data were obtained from 3 different samples for each developmental stage. Colors of dots indicate the normalized length.

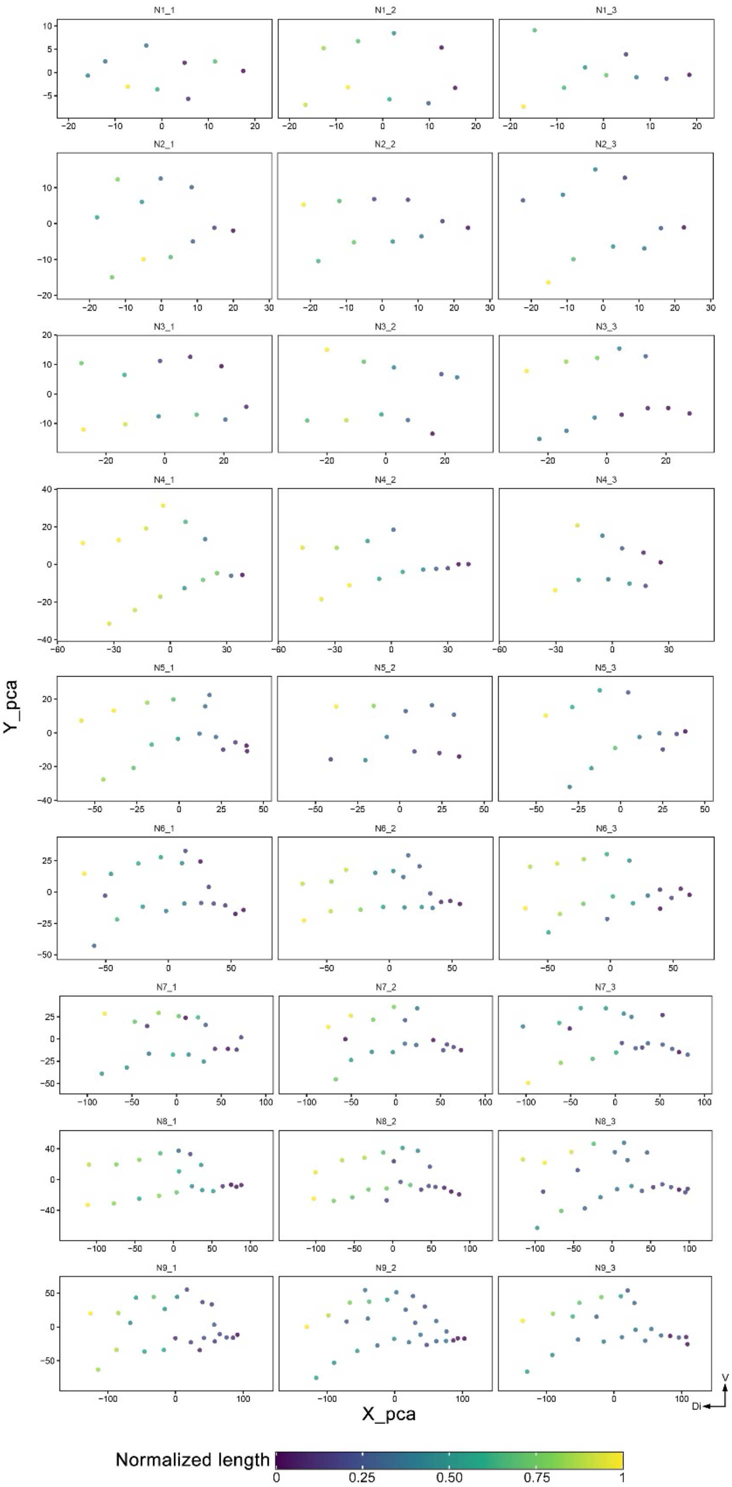


**Fig. S4  Standardized spatial distributions of lateral hair-plate sensilla across developmental stages.** Each dot represents an individual sensillum for the lateral hair plate of the antennal base, plotted in the standardized XY-plane after principal component analysis (PCA) alignment (See Material and Methods). The data were obtained from 3 different samples for each developmental stage. Colors of dots indicate the normalized length.
